# The human cytomegalovirus UL116 glycoprotein is a chaperone to control gH-based complexes levels on virions

**DOI:** 10.1101/2020.11.18.387126

**Authors:** Giacomo Vezzani, Diego Amendola, Dong Yu, Sumana Chandramuli, Elisabetta Frigimelica, Domenico Maione, Marcello Merola

## Abstract

Human cytomegalovirus (HCMV) relies in large part upon the viral membrane fusion glycoprotein B (gB) and two alternative gH/gL complexes, gH/gL/gO (Trimer) and the gH/gL/UL128/UL130/UL131A (Pentamer) to enter into cells. The relative amounts of the Trimer and Pentamer vary among HCMV strains and contribute to differences in cell tropism. Although the viral ER resident protein UL148 has been shown to interact with gH to facilitate gO incorporation, the mechanisms that favor the assembly and maturation of one complex over another remain poorly understood. HCMV virions also contain an alternative non-disulfide bound heterodimer comprised of gH and UL116 whose function remains unknown. Here, we show that disruption of HCMV gene *UL116* causes infectivity defects of ~10-fold relative to wild-type virus and leads to reduced expression of both gH/gL complexes in virions. Furthermore, gH that is not covalently bound to other viral glycoproteins, which are readily detected in wild-type HCMV virions, become undetectable in the absence of *UL116* suggesting that the gH/UL116 complex is abundant in virions. We find evidence that UL116 and UL148 interact during infection indicating that the two proteins might cooperate to regulate the abundance of HCMV gH complexes. Altogether, these results are consistent with a role of UL116 as a chaperone for gH during the assembly and maturation of gH complexes in infected cells.

## INTRODUCTION

Human cytomegalovirus (HCMV) infects most of the population primarily with an asymptomatic infection in immunocompetent individuals followed by a lifelong latent infection persisting in precursors of dendritic and myeloid cells (1–3). Reactivation and re-infection is a serious health problem in immunosuppressed patients where it represents the major causes of severe diseases or fatal outcome (4). In transplanted recipients, HCMV accelerates the rate of graft failure and vascular diseases (5). Furthermore, HCMV congenital infection remains a major problem associated to fetus neurodevelopmental delay or hearing/vision defects at birth (6).

This wide plethora of HCMV-associated disease likely relates to the ability of the virus to infect a diverse range of cell types, including epithelial and endothelial cells, fibroblasts, monocyte/macrophages, dendritic cells, hepatocytes, neurons, and leukocytes (7). This broad cell tropism may reflect the relative abundance of distinct glycoprotein complexes in the virion envelope. Together with glycoprotein B (gB), the gH/gL dimer comprises the “core membrane fusion machinery” conserved among all herpesviruses and is likely to regulate the fusogenic activity of gB (8). However, HCMV encodes a set of proteins that bind alternatively to gH/gL that modify or regulate the activity of the gB-gH/gL core fusion machinery leading to different tropism during the virus spreading in host cells (9). In particular, in HCMV, gH/gL exists on the viral surface as part of a trimeric complex with gO (gH/gL/gO, Trimer) or as a pentameric complex with UL128, UL130 and UL131A (gH/gL/UL128/UL130/UL131A, Pentamer) (10, 11). The presence of the Trimer, although required for entry into all cell types, is sufficient only for fibroblast infection while virus carrying Pentamer greatly expand cell tropism (12) recognizing different cellular receptors. Fibroblast entry relies on Trimer binding to platelet-derived growth factor receptor alpha (PDGFR-α) and ectopic expression of this receptor in PDGFR-α non-expressing cells restores infection of HCMV lacking Pentamer (13–15), As for the Pentamer, two groups have recently reported the identification of distinct receptors responsible for epithelial tropism. Using a cell-independent screening on purified ectodomain of single transmembrane human receptors, Martinez-Martin *et al.* have identified Neuropilin-2, that recognize the pUL128 and pUL131A subunit on the Pentamer, as an essential molecule for HCMV entry (16). Via a CRISPR/Cas9 genetic screening of human cells, E. *et al.* identified the 7 TM olfactory receptor OR14I1 associated with G proteins as Pentamer target required for endocytosis of the virus and subsequent infection (17).

While differential expression of these receptors influences permissivity to HCMV infection, at least for fibroblasts and epithelial cell types the viral infectivity relies upon the presence of the two gH-based complexes. The relative abundance of Trimer and Pentamer complexes in virions is strain-specific and influence cell tropism and infectivity (18). How the formation of the two gH/gL complexes is regulated at the molecular level remains currently largely unknown although two recent reports identified two potential players. Li *et al.* identified an ER-resident viral protein encoded by the UL148 gene (UL148) that influences the ratio of Trimer to Pentamer and the cellular tropism of HCMV virions (19). Deletion of UL148 from the viral genome impairs incorporation of the Trimer into virions, leading to a reduced capacity of viral particles to establish infection in fibroblasts while increasing level of infection in epithelial cells (19). The opposite outcome was observed by Luganini *et al.* using an *US16*-null virus which generated a Pentamer-deprived viral progeny that resulted unable to entry epithelial/endothelial cells (20). Whether these two viral proteins participate to form a tropism switch during the HCMV life cycle is unknown, however, this finding would imply a more complex system likely involving host proteins.

Among HMCV envelope proteins we identified UL116 as a gH interacting protein that forms noncovalent dimers alternative to gH/gL (21). In transient expression, gH and UL116 do not exit the ER unless they are co-expressed. The gH/UL116 complex migrates through the secretory pathway in the absence of other viral subunits suggesting that the antagonism with gL occurs once in the ER. Although the viral envelope localization of UL116 indicates a direct role in viral infection, its competition with any other gH/gL-based complex might reflect a role in the shaping of the envelope complexes.

In this work we found that, in the absence of UL116, cell-free viral spreading is reduced of about 10-folds with evidence of envelope gH/UL116 involvement. We addressed the role of UL116 in the early formation of gH-based viral envelope complexes and its interaction with UL148. We generated HCMV TR strain lacking the expression of either UL116 or UL148 to analyze the contribution of UL116 to complex choice and its potential interaction with UL148. We found that UL116 expression is required for the wt levels of both Trimer and Pentamer in virions produced by fibroblasts and epithelial cells. Furthermore, we also revealed a direct interaction of UL116 with UL148 in cells. These data collectively support the model of UL116 chaperoning gH during the early phases of complexes assembly.

## Materials and Methods

### Protein purification, reagents, plasmids and antibodies

Trimer, Pentamer and gH/UL116 heterodimer were purified as previously described (21, 22). Primary antibodies: anti-Pentamer was raised by immunizing rabbits with purified whole Pentamer protein (22)and purifying IgG from the serum over Protein A column. UL116 monoclonal antibody was produced in mouse following immunization with purified gH/UL116 and hybridomes screening. mouse mAb to Cytomegalovirus IE1 and IE2 (Abcam, ab53495), mouse mAb to Cytomegalovirus pp65 (Abcam, ab6503), rabbit pAb to Strep tag (Abcam, ab119810), 6xHis TagAntibody (Invitrogen, MA1-213115), anti-KDDDDK Tag antibody (Invitrogen, MA1-91878), Monoclonal antibody anti-GAPDH produced in mouse (SIGMA, G8795-200UL), Myc Tag monoclonal antibody (Sigma-Aldrich, 05-724), anti-gH human honoclonal antibody MSL109 was a generous gift of Dr. Adam Feire of the Novartis Institute for Biomedical Research (NIBR, Cambridge, MA, USA).

Secondary antibodies used are: Goat anti-mouse IgG (H+L) highly cross-adsorbed Alexa fluor plus 647 secondary antibody (Sigma-Aldrich, A32728), Goat anti-mouse IgG (H+L) secondary antibody HRP (Invitrogen, 62-6520) and Goat anti-rabbit IgG (H+L) secondary antibody HRP (Invitrogen, 62-6120).

HEK-293T transfections were carried out with Lipofectamine 2000 (Thermo Fischer) according to the manufacturer’s protocol. The HEK-293T transfected cells were trypsinized 48h post-transfection and treated for immunoprecipitation assays

gH_myc, UL116 and UL148-6XHIS were expressed following cloning of the codon optimized sequence in pcDNA3.1(−) plasmid.

All primers used are listed in Table 1.

**Table 1.**
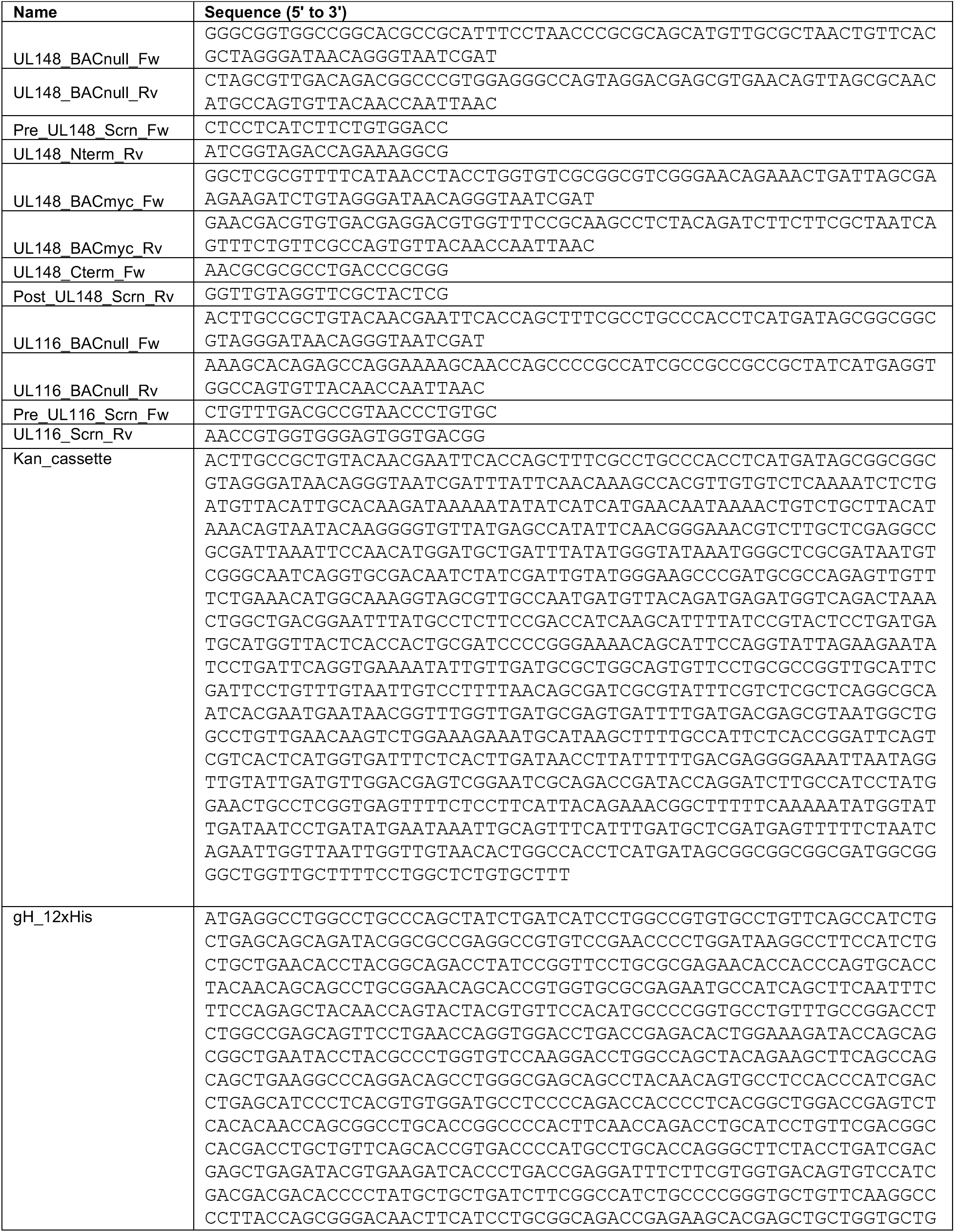
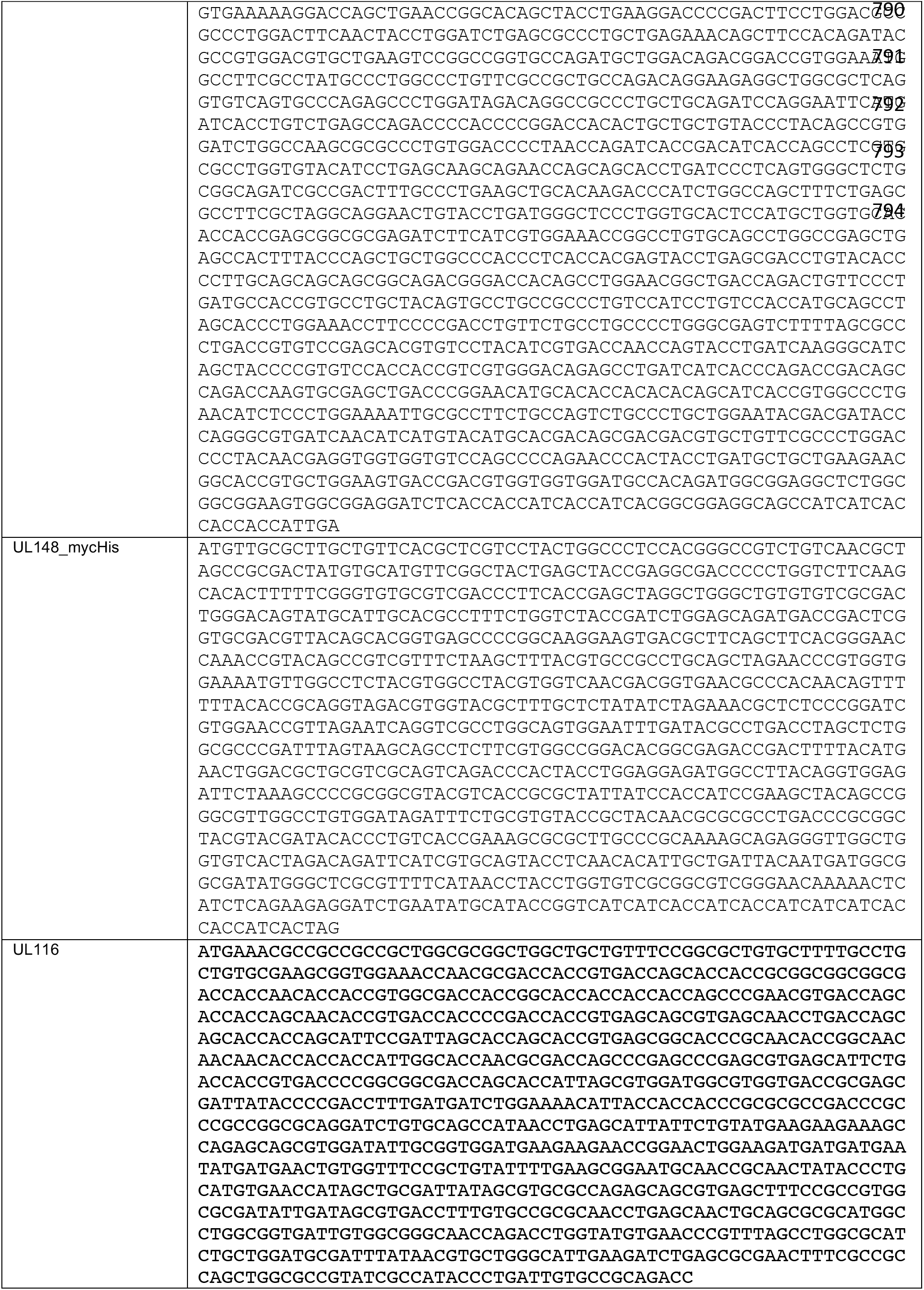
Oligonucleotides and synthetic DNA used in this study.

### Binding assay to HFF-1 and ARPE-19 cells

For the binding of gH/UL116, Trimer and Pentamer to cells, trypsinized HFF-1 or ARPE-19 were divided in identical aliquot of 3 × 10^5^ cells. Cells were first incubated for 20 min at room temperature (RT) with Live/Dead Aqua diluted 1:400 in PBS and then for 60 min with blocking buffer (PBS with 1% Bovine Serum Albumin (BSA) and 200 μg/ml of gH/UL116, Trimer or Pentamer recombinant complexes were incubated for 60 min at RT. All complexes were 6xHis-tagged. After three washes in PBS, mouse monoclonal anti-His and Alexa Fluor 647-conjugated anti-mouse antibodies were used to reveal the binding. A total of 10^5^ cells were analyzed for each histogram using FACS BD Canto II (Becton Dickinson, Heidelberg, Germany).

### Cell Lines

HFF-1 (Human [*Homo sapiens*] skin/foreskin normal fibroblasts; SCRC-1041), MRC-5 (Human [*Homo sapiens*] lung normal fibroblasts; CCL-171), ARPE-19 (Human [*Homo sapiens*] retinal pigmented normal epithelial cells; CRL-2302), HEK293T (Human [*Homo sapiens*] embryonic kidney epithelial cells; CRL-1573) cells were obtained from ATCC. HFF-1 cells were cultured in Dulbecco’s Modified Eagle Medium (DMEM, ATCC 30-2002) supplemented with 15% fetal bovine heat inactivated serum (FBS, ATCC 30-2020), 100 I.U./mL penicillin and 100 mg/mL streptomycin (Penicillin-Streptomycin, internally produced). MRC-5 and HEK293T cells were cultured in Eagle’s Minimum Exential Medium (EMEM, ATCC 30-2003) supplemented with 10% fetal bovine heat inactivated serum, 100 I.U./mL penicillin and 100 mg/mL streptomycin. ARPE-19 cells were cultured in Dulbecco’s Modified Eagle Medium/Nutrient Mixture F-12 (DMEM/F-12, ATCC 30-2006) supplemented with 10% fetal bovine heat inactivated serum, 100 I.U./mL penicillin and 100 mg/mL streptomycin. All cell lines were grown at 37°C with 5% CO_2_.

### Viruses

A bacterial artificial chromosome (BAC) containing the genome of the HCMV TR strain was obtained from Oregon Health Science University (23) and was integrated with a GFP immediate early expressing gene cassette in the intergenic region between US32 e US33A genes. TR, a clinical HCMV strain derived from an ocular vitreous fluid sample from a patient with HIV disease (24), was cloned into a BAC after limited passage in fibroblasts (23). HCMV strain TR-GFP (TRG) and each recombinant virus were propagated in HFF-1 fibroblasts grown to 70-80% confluency, as previously described (Cell Lines, STAR Methods), using infectious supernatants at a MOI of 1. Infection of ARPE-19 cells was performed at a MOI of 5. Infection was visualazed at 24 hpi (hours post-infection) by GFP-fluorescence inside cells. At 100% CPE (or GFP signal) or 50% of cells detached from the plate, medium supernatant was collected and cleared of cell debris by centrifugation for 15 min at 4,000 × g at 4°C before aliquoting and storing at −80°C.

To titrate viruses, we used a Titration Assay previously described (25) with minor modifications. In brief, 5-fold serial dilutions of samples were performed in DMEM supplemented with 1% fetal bovine heat inactivated serum and 1 mM sodium pyruvate, and 150 μl of each dilution was applied to duplicate wells of a 96-well flat bottom cluster plate containing 2 × 10^4^ HFF-1 fibroblasts, incubated over-night (O/N) at 37°C with 5% CO_2_ before infection. At 24 hpi, the infected cells were trypsinized and transferred in a 96-well round bottom cluster plate. To evaluate the number of cells with GFP-signal, we performed FACS analysis with BD LRSII Special Order System (Becton Dickinson, San Jose, CA) equipped with High Throughput Sampler (HTS) option. Titer was calculated using the following equation: Titer (IU/ml) = (N × P)/(V × D) [Note: N = Cell Number in each well used for infection day; P = percentage of GFP positive cells (considering the dilution virus exhibiting GFP signal < 40%); V = virus volume used for infection in each well (ml); D = dilution fold; IU = infectious unit].

For kifunesine treatment, six T75 flasks were seeded with HFF-1 cells and infected with HCMV TRG (2 flask) and TRG-*UL116*-null (2 flasks) at MOI of 1. After 72 hours, kifunesine (Sigma-Aldrich, K1140) was added to the final concentration of 5 μM in the culture media of three flasks (uninfected HFF-1, TRG infected HFF-1 and TRG-*UL116*null infected HFF-1). The same amount of sterile distilled water was added to the remaining 3 flasks. 48 hours after drug treatment, cells were harvested, lysed and treated for western blot analysis in reducing and nonreducing conditions.

### BAC Mutagenesis

To generate recombinant viruses a Two-step Red-mediated recombination method has been used as previously described (26) with minor modifications. BAC TR-GFP was used as starting template. In brief, kanamycin resistance cassette, flanked by I-SceI restriction enzyme cleavage sites, was amplified from pEPkan-S shuttle vector using primers containing homologous regions for the integration in the region of interest. Recombination events were performed with *E. coli* GS1783 strain containing a BAC clone of the HCMV TR-GFP (TRG) strain, the lambda Red system under the control of a heat-inducible promoter and the I-SceI genes under the control of an arabinose-inducible promoter (27). The first recombination step consists in the electroporation of the purified PCR-amplified cassette in competent, heat-induced GS1783 cells. Positive clones for cassette integration were selected based on kanamycin resistance and screened both by PCR and sequencing. The second recombination was triggered through both heat-shock and arabinose and results in the excision of the kanamycin resistance, leaving the mutation in frame with the gene of interest. Putative clones were screened by PCR and sequencing analyzed by Vector NTI.

### Reconstitution of infectious viruses

To reconstitute the virus MRC-5 fibroblasts were electroporated (nucleofected) using a Cell Line Nucleofector Kit V Lonza VCA-1003) according to the manufacturer’s protocol. In brief, for each reaction, 1 × 10^6^ freshly trypsinized MRC-5 fibroblasts were pelleted by centrifugation at 300 × g for 5 min, washed two times with PBS and then resuspended in a solution containing 1,5 μg of BAC and 0,3 μg of pcDNA3.1-pp71 plasmid premixed with 100 μL of Nucleofector solution (82 μL of Nucleofector solution and 18 μL of supplement). Cotransfection of HCMV protein pp71-expressing plasmid markedly increases the efficiency of virus reconstitution from transfection of infectious viral DNA since pp71 acts as a viral transactivator to help initiate lytic infection (28). The cell suspension was then electroportated using a Nucleofector II (program D-023) and then plated and cultured in DMEM supplemented with 1% fetal bovine heat inactivated serum and 1 mM sodium pyruvate. 24h after electroporation, medium was changed and cells were cultured by standard methods. When cells exhibited 100% CPE (or GFP signal, observed with a Zeiss Axiovert 200) or 50% of cells were detached from the plate, medium supernatant was collected and cleared of cell debris by centrifugation for 15 min at 4,000 × g, 4°C before aliquoting and storing at −80°C. To determine virus titer the “Titration Assay” has been performed as previously described (Viruses, STAR METHODS).

### HCMV Virions purification

The supernatant of infected cells was collected 7 days (HFF-1) or 8 days (ARPE-19) after infection and centrifuged for 15 min at 4,000 × g, 20°C to clear all cell debris. Cleared supernatant was transferred to polycarbonate ultracentrifuge tubes under lied with 20% sucrose cushion and centrifuged at 30,000 rpm in a Beckman SW32Ti rotor for 50 minutes. The virus-containing pellet was solubilized in 1% Triton X-100 in PBS and protease inhibitors (EDTA-free EASYpack Protease Inhibitor Cocktail (Sigma-Aldrich) and finally equilibrated in SDS-PAGE loading buffer for western blot analysis.

### Immunoprecipitations

HFF-1 cells were infected at MOI of 1 with HCMV TRG-wt, TRG-UL148-myc or TRG-*UL148*-null. Infection was allowed to proceed for 6 DPI and then cells were washed in 1× PBS, lysed with Mammamlian CelLytic (Sigma-Aldrich) in presence of protease inhibitors. Five hundred micrograms of total protein extracts were incubated overnight at 4°C with 5 μg of Myc Tag Monoclonal Antibody, anti-UL116 Monoclonal Antibody or anti-gH Human Monoclonal Antibody. Complexes were pulled down using Dynabeads Protein A/G (Sigma-Aldrich, 14321D) according to the manufacturer’s protocol. Recovered beads were washed in lysis buffer and then boiled for 5 min in 2X SDS-PAGE loading buffer with reducing agent. Eluted proteins were separated on SDS-PAGE and immunoblotting performed as described above.

A similar procedure was applied to recover immunocomplexes from transfected HEK293T cells. 3 × 10^5^ HEK293T cells per well were seeded in a 6 wells plate and incubated O/N at 37°C. The day after, cells were transfected with 10 μg of each plasmid. Extracts were then used for immunoprecipitation procedure using 5 μg of each antibody (gH Human Monoclonal Antibody, myc Tag Monoclonal Antibody and UL116 H4).

### Immunoblotting

Proteins were separated by sodium dodecylsulfate-polyacrylamide gel electrophoresis (SDS-PAGE) on 4-12% polyacrylamide pre-cast gels (Bolt 4-12% Bis-Tris Plus Gels) under reducing or nonreducing conditions. Proteins were transferred to nitrocellulose membranes (iBlot 7-Minute Blotting System, Invitrogen), and membranes were blocked with PBS containing 0.1% Tween 20 (ThermoFisher, TA-125-TW) and 10% powdered milk (Sigma-Aldrich, M7409). Antibodies were diluted in PBS containing 0.1% Tween 20 and 1% powdered milk. For detection of primary antibody binding, horseradish peroxidase-conjugated anti-rabbit or anti-mouse IgG antibodies and the Chemiluminescent Peroxidase Substrate (Sigma-Aldrich, 34578) were used, according to the manufacturer’s instructions. The densitometric analysis of signal intensity in Western blotting was performed with ImageLab software.

## RESULTS

### Construction of HCM TR-GFP (TRG) strain mutants and cell-free viral growth in human fibroblast and epithelial cells

To characterize the role of UL116 in viral pathogenesis, we first checked cell-free infectivity in the absence of UL116. We generated a recombinant virus that do not express the UL116 protein by inserting a stop codon close to the N-terminus of the UL116 open reading frame (ORF). UL148 is a HCMV ER resident protein reported to bind gH and influence the gH-based complexes formation (19). We constructed a mutant virus lacking UL148 expression to be studied in parallel. Finally, to detect UL148 in infection, we constructed a recombinant virus expressing UL148-myc tagged protein. In figure 1 is depicted the map of viruses used in this study. All viruses were generated from the bacterial artificial chromosome (BAC) containing the HCMV TR strain to which the GFP gene was introduced between US32 e US33A genes. This template was used to generate recombinant viral genomes via a marker-less two-step RED-GAM BAC mutagenesis (29). The TR-GFP wt (to which we will refer to as TRG) was used to generate the TRG-*UL116-*null, the mutant lacking UL116 expression by insertion of a single nucleotide between residues 4-5 to generate stop codon immediately afterwards, and the TRG-UL148-myc, containing the tag at the C-terminus. The latter was used as template to generate the TRG-*UL148-* null in which a stop codon was introduced at position 4 of the *UL148* ORF.

**Figure 1.**
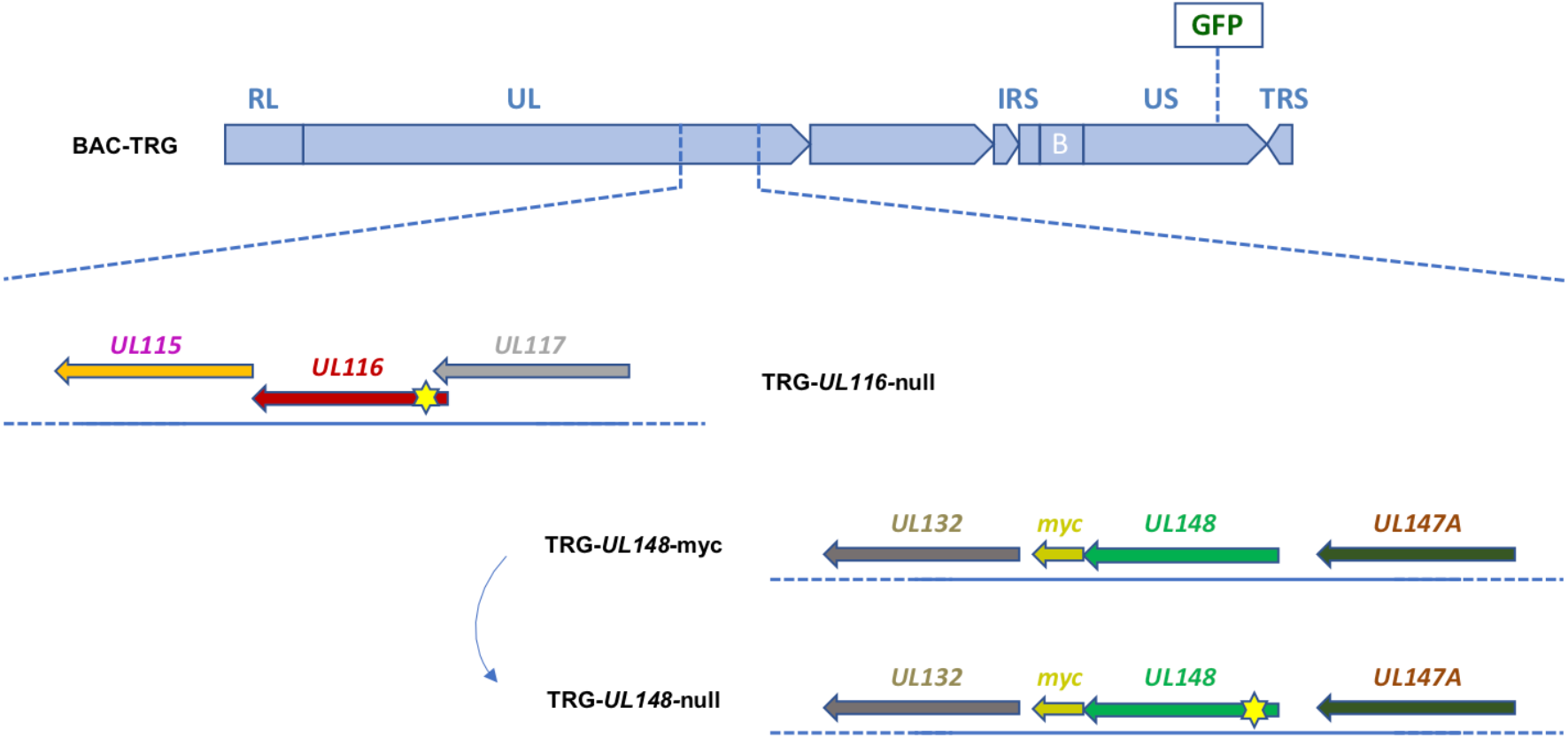
Schematic representation of recombinant HCMV mutants generated. Each mutant is generated through the BAC Mutagenesis Technique. The backbone sequence, common for each mutant shown, consists of the HCMV TR-wt strain cloned into a BAC and containing a GFP CDS insertion, under control of an Immediate Early CMV promoter, in an intergenic region between US32 e US33A genes (BAC-TRG). The TRG-*UL116-*null clone was generated using the BAC-TRG as template with a single nucleotide insertion in the CDS (between nucleotides in position 4-5) of the *UL116* gene causing a frameshift and a STOP codon formation. The TRG-UL148-myc clone was generated using the BAC-TRG as template inserting the sequence encoding for a myc-tag in frame at Carboxy terminal. The TRG-*UL148*-null clone was generated from the BAC-TRG-UL148-myc by mutation of the codon at position 4 into a STOP codon. Reconstitutions of the infectious viruses was performed as detailed in M&M. Yellow stars indicate the approximative position of the stop codon insertion.

Fibroblasts have always been the standard cell type for isolation and propagation of HCMV from patient samples and are still the most efficient producer cell line irrespective of the virus strain. As first, we investigated cell-free replication into human foreskin fibroblasts (HFF) to verify if the mutations introduced in our recombinant viruses could have effect on viral growth. HFF-1 cells were infected at a multiplicity of infection (MOI) 1 and aliquots of media collected up to 7 days. Production of cell-free virus was measured by titrating infectious viruses secreted in cell media on fresh HFF-1 cells. As shown in figure 2A, replication of the TR-*UL148-*null virus was identical to the wt while the TRG-*UL116-*null show a reduction of about 10 times. These data indicate that eradication of UL116 expression influence viral replication and/or infectious ability.

**Figure 2.**
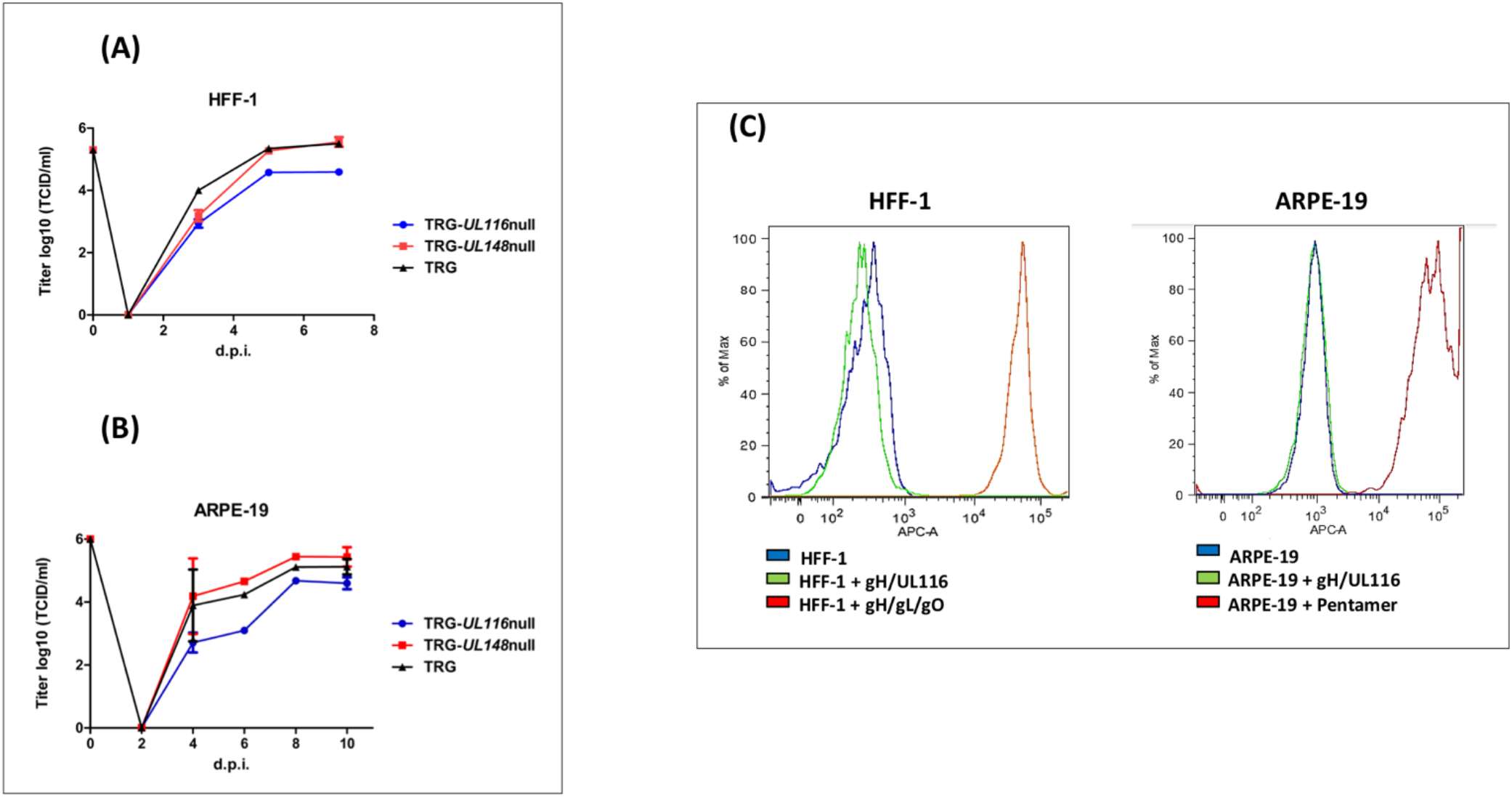
Growth curve of TRG, TRG-UL116null and TRG-UL148null and binding of recombinant gH/UL116 to HFF-1 and ARPE-19. **A**) HFF-1 cells were infected at MOI 1 and cultured for 7 days. At the indicated times, aliquot of the medium were withdrawal and viral titer assessed on fibroblasts. **B**) ARPE-19 cells were infected at MOI 5 and cultured for 8 days. At the indicated times, aliquot of the medium were withdrawal and viral titer assessed on fibroblasts. **C**) 200 μg/ml of recombinant gH/UL116, Trimer and Pentamer were incubated for 1 hr with 10^5^ cells in blocking buffer (PBS with 1% BSA). His tag was present on recombinant gH. HIS-Tag Monoclonal Antibody [HIS.H8] and then Alexa Fluor 647-conjugated anti-mouse secondary antibody were used to reveal complexes bound to cells. A total of 10^5^ cells were analyzed for each histogram using FACS BD Canto II (Becton Dickinson, Heidelberg, Germany).

Apart from fibroblasts, epithelial cells are one of the major targets of HCMV infection and are assumed to play an important role during host-to-host transmission since they lay all external body surfaces. We sought to repeat the same analysis on ARPE-19 epithelial cells to verify if mutants had a differential tropism. ARPE-19 cells were infected at a multiplicity of infection (MOI) 5 and aliquots of media collected up to 10 days. As it can be seen in figure 2B, viral secretion from epithelial cells displayed one day delay in viral secretion compared to fibroblasts but the titers measured at plateau mirrored what observed in fibroblasts. The TRG-*UL116-*null virus showed 0.65 log lower titer at plateau with respect to the wt or TRG-*UL148-*null. To note that our TR strain not expressing UL148 did not reproduced the behavior reported by Li et al. who found and increased epithelial tropism (19).

Results from these experiments suggest that the TR strain impaired cell-free virus production in absence of UL116 is cell type independent.

### Soluble gH/UL116 does not bind to fibroblasts and epithelial cells

We speculated that virion envelope-bound gH/ UL116 dimer might facilitate virus to attach the host cell contributing to viral attachment and/or entry, and therefore lacking UL116 might be responsible for the observed reduction in viral titer of the TRG-*UL116*-null virus. To test this hypothesis, we investigated the binding of the gH/UL116 heterodimer to fibroblasts and epithelial cells. We expressed and purified soluble recombinant gH/UL116 tagged with 6xHis and strep respectively (21) and checked binding to HFF and ARPE-19 cells by FACS analysis. Recombinant Trimer and Pentamer were used as control. As expected, strong binding of the Trimer to fibroblasts and of the Pentamer to epithelial cells were revealed whereas no binding of gH/UL116 to both cells could be revealed (figure 2C). This finding indicates that gH/UL116 does not target a high affinity receptor on cultured fibroblast and epithelial cells.

### Expression levels of the major HCMV envelope proteins in infected cells and their incorporation into secreted virions

Zhang et al. asserted that the differential tropism and infectivity of distinct strains is also function of the relative levels of Trimer, Pentamer and gH/gL carried by virons (18). This observation encouraged us to verify the levels of expression of the gH-based complexes proteins in absence of UL116 and/or UL148 both in virions and in infected cells. HFF-1 cells were infected with wt and mutant viruses at MOI of 1 for 7 days, then culture media were collected for viral purification on sucrose cushion gradient and cells were harvested. Pelleted virions and cells were then lysed in detergent containing buffer and analyzed by western blot in both reducing and nonreducing conditions. Free gH, Trimer, and Pentamer complexes were analyzed on SDS-PAGE in non-reducing conditions in which free gH, gH/gL/gO (Trimer), and gH/gL/UL128 (Pentamer) complex migrated at an apparent MWs around 85-90, 260, and 150 kDa respectively (Figure 3) (22). Densitometric analysis of the immunoblot were performed and the intensity of the bands corresponding to the different species was compared.

**Figure 3.**
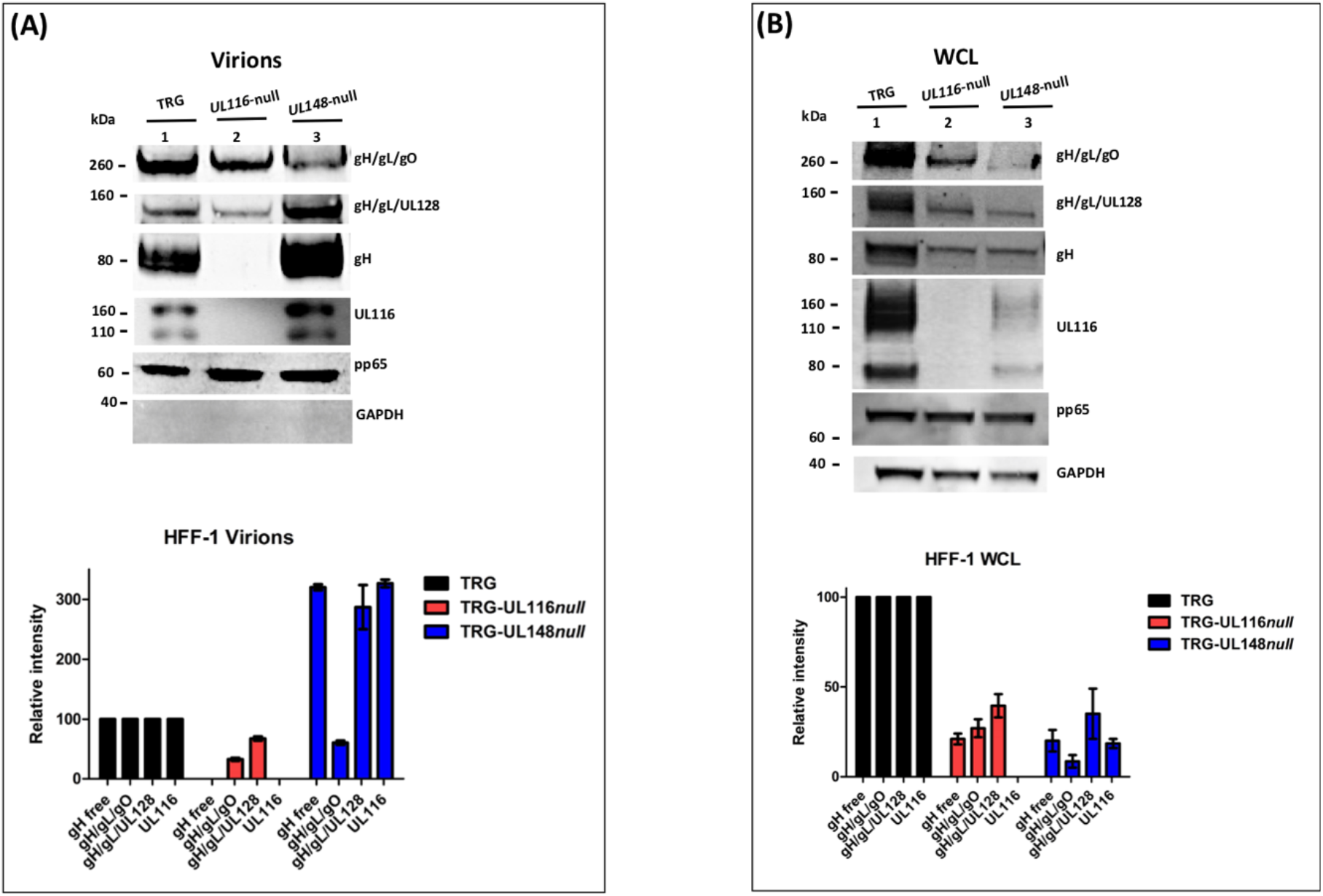
Loss of *UL116* or *UL148* gene products alters the ratio of gH/gL complexes in HFF-1 cellular extracts and virions. UL116 and GAPDH were revealed in reducing conditions. gH, gH/gL/gO, gH/gL/UL128 were probed with anti-Pentamer polyclonal antibodies in nonreducing conditions. Western blots showed are representative of at least three independent experiments. On the bottom, densitometric analysis of the corresponding immunoblot are shown. Densitometric values of complexes present in the wt virus are considered 100%. The standard deviation indicated in graphs were obtained by the densitometric values from three independent experiments and normalized on the intensity of the pp65 viral marker. Viruses used for infection are indicated on the top of each lane. **A**) Viral pellets from a 75 cm^2^ flasks for each sample were lysed in 50 μl of 1% Triton X-110 in PBS. 5 μl aliquots were runned in SDS-PAGE in reducing conditions, transfered to a nitrocellulose membrane and probed with anti-pp65 whose levels were used to normalized the consecutive loads. Two aliquots of each sample were then loaded on 4-12% PAGE-SDS in reducing and nonreducing conditions respectively and treated for immunoblotting. Virus lysates are indicated on the top of each lane **B**) Equal amount of total proteins (BCA) from 7 DPI whole cell lysates (WCL) of HFF-1 were separated (NuPage, Invitrogen) and treated for western blot analysis.

gH-based complexes immunoblot on virions is shown in figure 3A. The TRG wt showed Trimer as major complex on the envelope as previously described (18) and high level of gH although only a minority as gH/gL. The TRG-*UL116*-null mutant revealed a strong reduction of Trimer almost comparable to the expected band found in the TRG-*UL148*-null virions (lanes 2 and 3 of figure 3A). The levels of Pentamer carried by the two mutated viral particles showed opposite outcome. TRG-*UL148*-null virions exhibited higher levels of Pentamer (lane 3 of figure 3A) as previously reported (19), whereas, in absence of UL116, considerable reduction of the Pentamer was observed (lane 2 of figure 3A). Interestingly, in TRG-*UL116*-null mutant virions, the levels of non-disulfide bound gH became undetectable. This suggests that a relevant amount of the viral gH not engaged in Trimer or Pentamer is normally present on the viral envelope associated to UL116. Finally, the TRG-*UL148*-null virions carried remarkably high levels of gH and UL116, likely as dimer, compared to the wt (lanes 1 and 2 of figure 3A). Altogether, these data indicate that the absence of UL116 impair incorporation of both gH/gL complexes in the viral particles whereas the loss of UL148 promotes increased incorporation in virions not only of Pentamer but also of the gH/UL116 dimer.

A representative western blot analysis of infected HFF-1 whole cell lysates (WCL) is displayed in figure 3B. Densitometric analysis consider values variation of three independent experiments normalizing on the intensity of the tegument protein pp65. Both mutants showed a reduced level of the Trimer and the Pentamer complexes (figure 3B). Cellular pool of free gH in the two mutants were reduced compared to the wt in an almost identical manner and not completely absent from the TRG-*UL116*-null mutant as observed in virions (compare lanes 2 of figure 3A and 3B). This suggests that contemporary expression of both UL116 and UL148 is required to completely stabilize intracellular pool of gL-free gH. Viral protein expressed in the cellular extracts of TRG-*UL148*-null mutant infected HFF-1 showed both a pronounced reduction of the Trimer and high levels of UL116. These results suggest a close relationship between gH, UL116 and UL148 in the ER of fibroblasts.

Identical experiments were performed on extracts from infected ARPE-19 cells and virions produced in this cell line. ARPE-19 cells were infected at a MOI of 5 and incubated for 8 days. At the end of this period, cell culture media were used for virus preparations on sucrose cushion while cells were harvested and lysed in detergent containing buffer. Representative western blots from these experiments are shown in figure 4 as well as densitometric analysis of the gH-based complexes, mediated on three independent experiments, are graphed in the bottom of the figures. The absence of UL116 lead to the disappearance of free gH on virions in nonreducing conditions (lane 2 of figure 4A) whereas we observed a roughly 50% reduction in infected cells (lane 2 of figure 4B). These results are consistent with the one obtained in fibroblasts (lanes 2 of figure 3A–4A and 3B-4B respectively) and with a role of UL116 in stabilizing and promoting gL-free gH incorporation into virions. The levels of the gH-based complexes carried by TRG-*UL116*-null virions produced by epithelial cells was less than half of both Trimer and Pentamer with respect to the wt (lanes 2 of figure 4A). As expected, TRG-*UL148*-null virions showed reduced Trimer and increased Pentamer but also higher amount of UL116 (lane 1 of figure 4A). Thus, unbalanced viral incorporation of gH-based complexes was observed in virion produced after infection of both cell lines and for both mutants.

**Figure 4.**
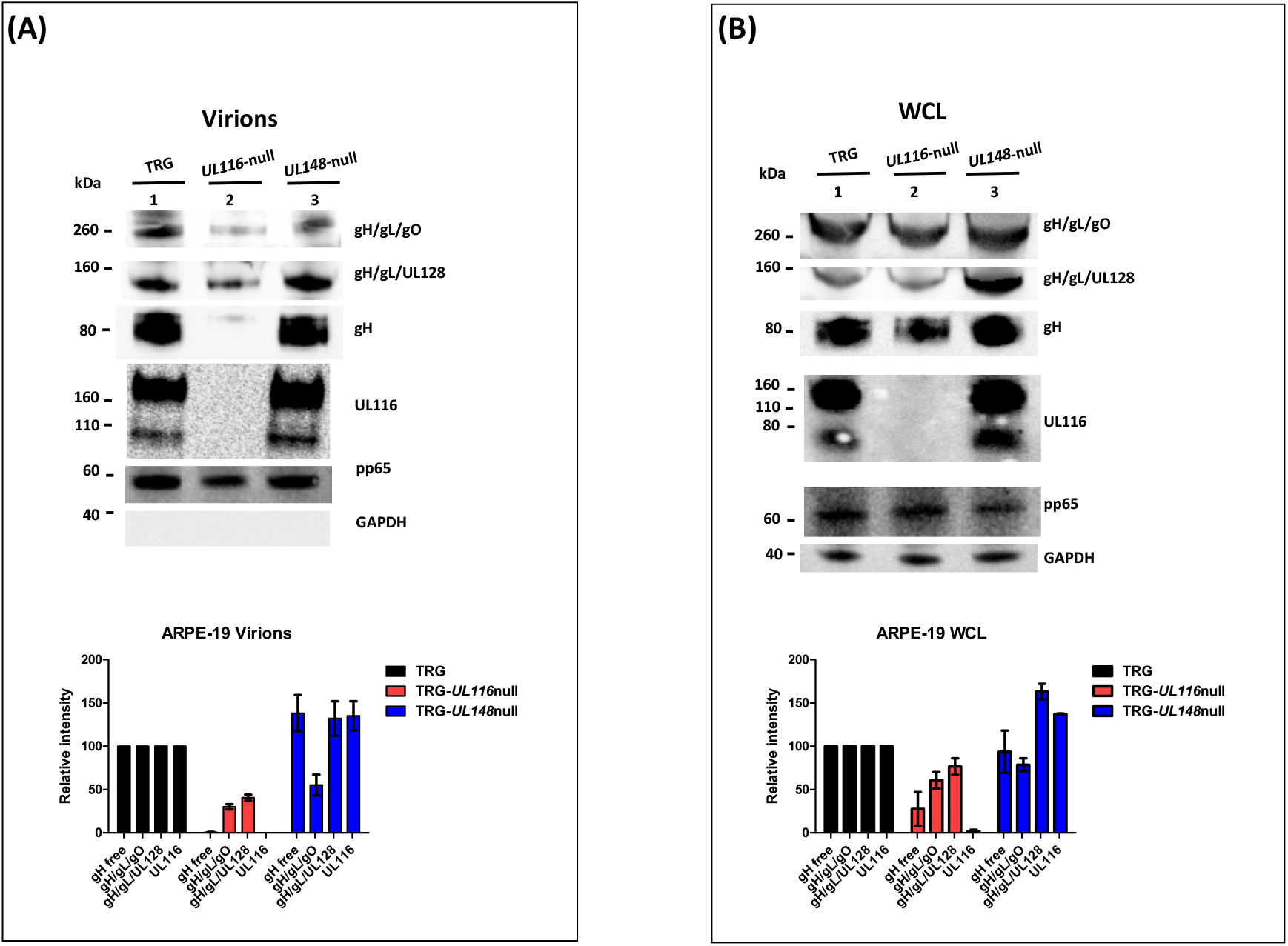
Loss of *UL116* or *UL148* gene products alters the ratio of gH/gL complexes in ARPE-19 cellular extracts and virions. Analysis was performed as specified in the legend of figure 3. Viruses used for infection are indicated on the top of each lane. **A**) Immunoblots of lysates of sucrose cushion-purified virions (8 DPI) **B**) Equal amount of total proteins (quantified by BCA) from 8 DPI whole cell lysates (WCL) of ARPE-19 infected with the viruses indicated on the top of the figure were separated on 4-12% PAGE-SDS (NuPage, Invitrogen) and treated for western blot analysis.

Analysis of the relative levels of gH-based complexes performed in cell lysates from wt and mutants infected ARPE-19 is shown in figure 4B. TRG-*UL116*-null mutant showed reduced levels of Trimer and Pentamer (lane 2 of figure 4B) although this reduction was less pronounced with respect to what observed in fibroblasts (graphs of figure 3B and 4B). Non covalently bound gH is present at low levels, the majority likely degraded for the absence of UL116. The levels of HCMV complexes in ARPE-19 cells infected with the TRG-*UL148*-null mutant differed from what found in fibroblasts. The levels of Trimer were almost as the wt indicating an intracellular accumulation without productive insertion into virions. The intracellular amount of non-covalently bound gH was equal to the wt (lane 3, figure 4B) suggesting that ARPE-19 may present factors that stabilize this glycoprotein that are absent in fibroblasts.

Taken together, this analysis reveals similar picture in virion compositions of particles derived from fibroblasts and epithelial cells assessing a crucial role for UL116 and UL148 to generate a pattern of gH complexes typical of the TR HCMV strain. Difference in the intracellular population of HCMV glycoproteins among the two different cell type could suggest differential pattern of interactors that, in epithelial cells, can stabilize HCMV species but not allow insertion on viral particles. Thus, both proteins may act in increasing proper assembly of gH-based complexes.

### Kifunesine treatment partially restore gH levels in TRG-UL116-null mutant

Data shown so far suggest that UL116 acts as gH “escort” protein implying that in its absence gH would be degraded faster by the endoplasmic reticulum associated degradation (ERAD) machinery. To test this hypothesis, we used the ER mannosidase inhibitor kifunesine that hinder mannose trimming of the oligosaccharide chain and further recognition by the ERAD factors (30). We reasoned that, in presence of this inhibitor, gH must accumulate in the ER and we analyzed extracts of infected cells treated and untreated with this drug. Results are shown in figure 5. In reducing conditions, gH levels in the cellular extract of TRG-*UL116*-null infected HFF-1 dropped to about 40% compared to the TRG wt (lanes 1 and 3 of figure 5) whereas was rescued to about 80% following kifunesine treatment (figure 5, lanes 2 and 4). As expected, in nonreducing conditions the base levels of free-gH in the extract derived from TRG-*UL116*-null infection dropped to 20% (figure 5, lane 3) and it was rescued to 40% upon kefunesine treatment. Level of two other HCMV proteins, gB and pp65, were not modified by the drug (figure 5). From this data we deduced that UL116 protects gH from an accelerated degradation.

**Figure 5.**
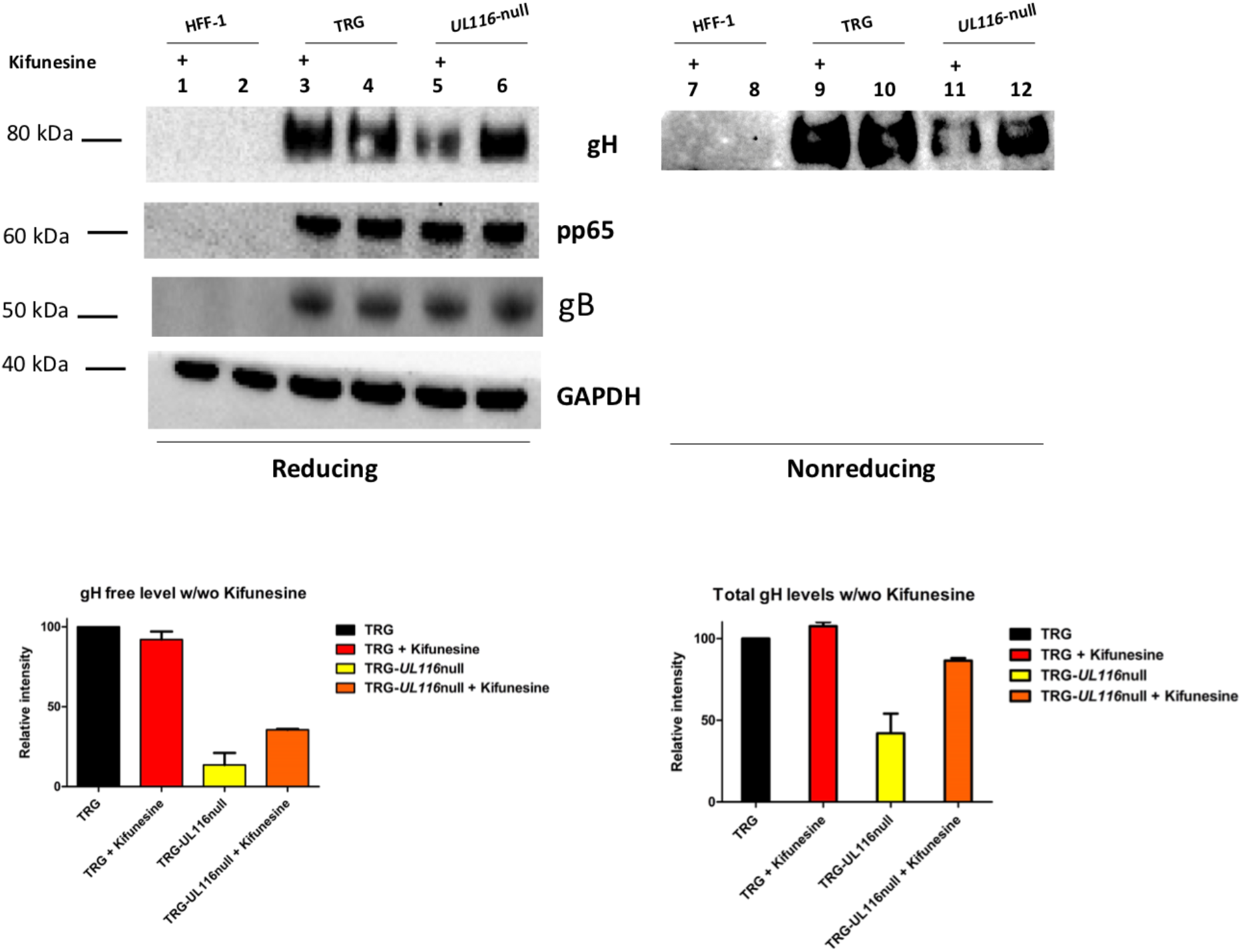
Free-gH levels in WCL from HFF cells infected with TRG and TRG-*UL116*-null under kifunesine treatment. HFF-1 cells were infected in duplicate with TRG wt and TRG-*UL116*-null and incubated 72 hrs. At the end of this period, 5 μM Kifunesine was added to one of the pair cultures and incubated for additional 48 hrs. Cells were then harvested, lysed and WCLs were treated for western blot analysis in both reducing (lanes 1-6) and nonreducing conditions (lanes 7-12). Non infected cells were treated identically and used as control.Protein separation was achieved on 4-12% NuSieve gels (Invitrogen) using equal amount of total protein (BCA). Rabbit polyclonal anti-Trimer was used to detect gH. Intensity of the GAPDH bands were used to normalize values reported on the graph.

### Co-immunoprecipitation in infected and transfected cells

Our data suggested that UL116 assist gH in its folding indicating a possible interaction with the viral ER resident protein UL148 which was shown to play a role in gH-based complexes choice. We asked if the two proteins could have a direct contact in the early stages of viral glycoproteins assemble. To this aim, we performed co-immunoprecipitation experiments in extracts from HFF-1 cells infected with TRG, TRG-UL148-myc and TRG-*UL148*-null. In absence of a specific anti-UL148 antibody we used a virus carrying myc-tag at the C-terminus of UL148. Immunoprecipitated samples were resolved in SDS PAGE and immunoblotted with specific antibodies. Expression of all individual proteins was revealed by western blot of non-immunoprecipitated WCLs (figure 6, lanes 14-16). In TRG and TRG-*UL148*-null mutant, no UL148 could be detected by the anti-myc antibody and only co-immunoprecipitation of gH by anti-UL116 (figure 6, lanes 7 and 11) and of UL-116 by anti-gH (figure 6, lanes 6 and 10) could be observed. In extracts infected with the TRG-UL148myc, however, UL148 was co-immunoprecipitated not only by anti-gH but also by anti-UL116 (figure 6, lanes 2 and 3). Although this result suggests a direct interaction between UL116 and UL148, it does not discriminate whether they interact directly or if they are simultaneously associated to the same protein such as gH that has been reported to bind independently with both proteins (19, 21). Our result is consistent with these reports since anti-gH antibodies co-immunoprecipitated both UL116 and UL148 (figure 6, lanes 2 and 6).

**Figure 6.**
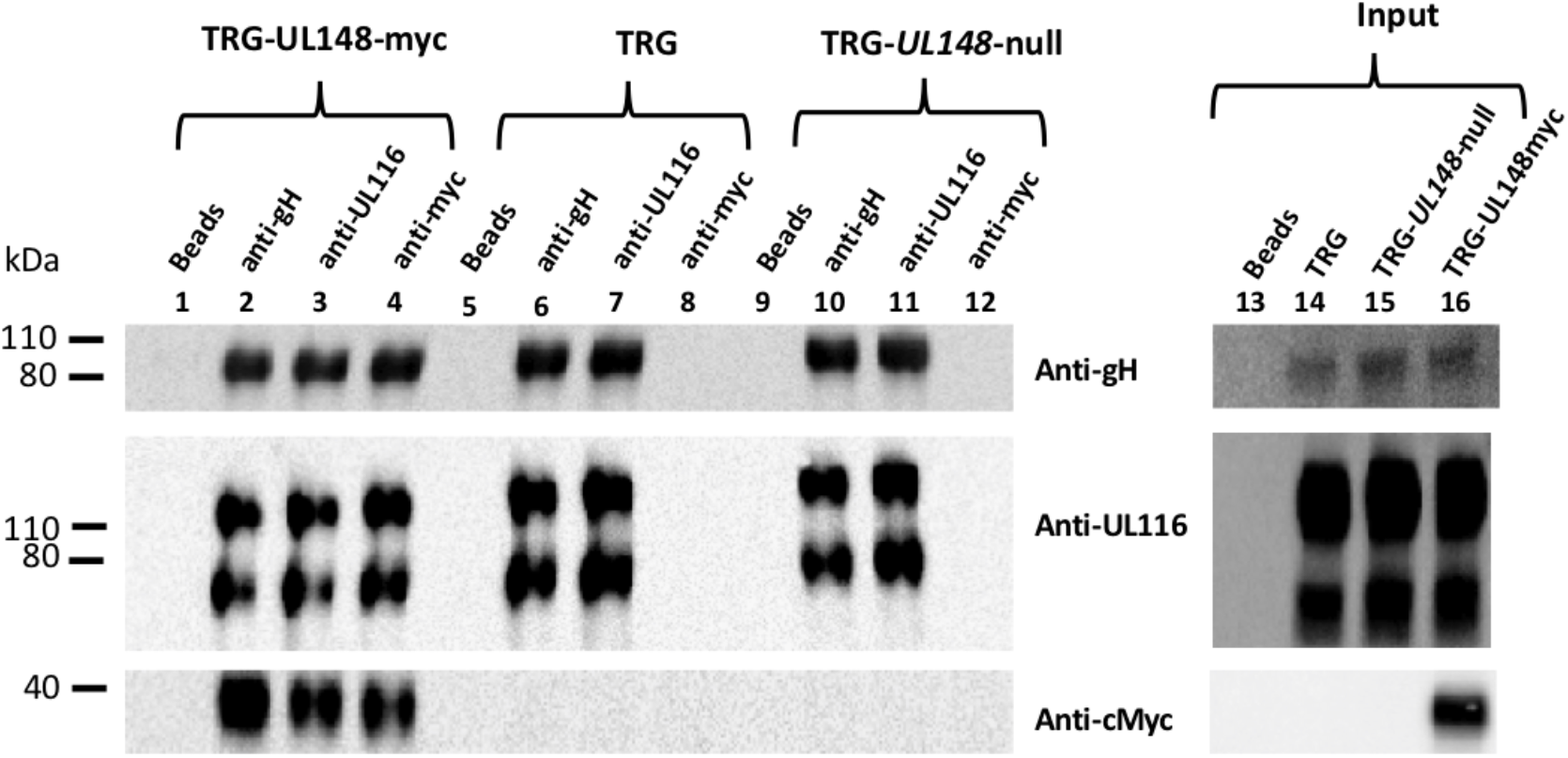
Co-immunoprecipitations of WCL from HFF cells infected with TRG, TRG-UL148myc and TRG-UL*148*-null. HFF-1 cells were infected for 6-days at MOI 1 with TRG, TRG-UL148myc and TRG-*UL148-*null. Infected and non-infected cell lysates were immunoprecipitated with beads only (lanes 1, 5 and 9), Anti-UL116 (H4) (lanes 2, 6 and 10), anti-myc (lanes 3, 7 and 11) and anti-gH (MSL-109) (lanes 4, 8 and 12) as indicated on the top of the lanes. Protein separation was achieved on 4-12% NuSieve gels (Invitrogen) and probed in western blot by anti-His, anti-myc and mAb F11 for gH, UL148 and UL116 respectively (indicated on the right). GAPDH was used as marker to normalize lysate amount and to exclude contamination in Immunoprecipitations. Input lysates (lanes 13-16) were also probed for detection of the individual HCMV proteins.

To provide evidence that UL116 and UL148 have a direct interaction, we performed co-immunoprecipitation in HEK-293T cells co-transfected with expression vectors for tagged individual HCMV proteins. For instance, plasmids used for transient transfection expressed 6xHis tagged gH, His/myc-tagged UL148 and Strep-tagged UL116. HEK-293T cells were transfected with different combination of the three expression plasmids and protein immunoprecipitations in whole cell lysates were carried out with human mAb MSL109, mouse mAb F11 and rabbit anti-myc for gH, UL116 and UL148 respectively. Proteins were revealed in western blot by anti-His, anti-myc, and mAb F11 for gH, UL148, and UL116 respectively. Figure 7 shows a representative result of such analysis. Expression of each protein was verified in western blot by immunoblotting an aliquot of WCLs (figure 7, lanes 17-24). As expected, both UL116 and UL148 co-immunoprecipitated with anti-gH (figure 7, lanes 2 and 3). Co-immunoprecipitation of UL148, via anti-myc antibody, pulled down both gH and UL116 glycoproteins when these species were individually co-expressed with UL148 (figure 7, lane 11 and 12 respectively).

**Figure 7.**
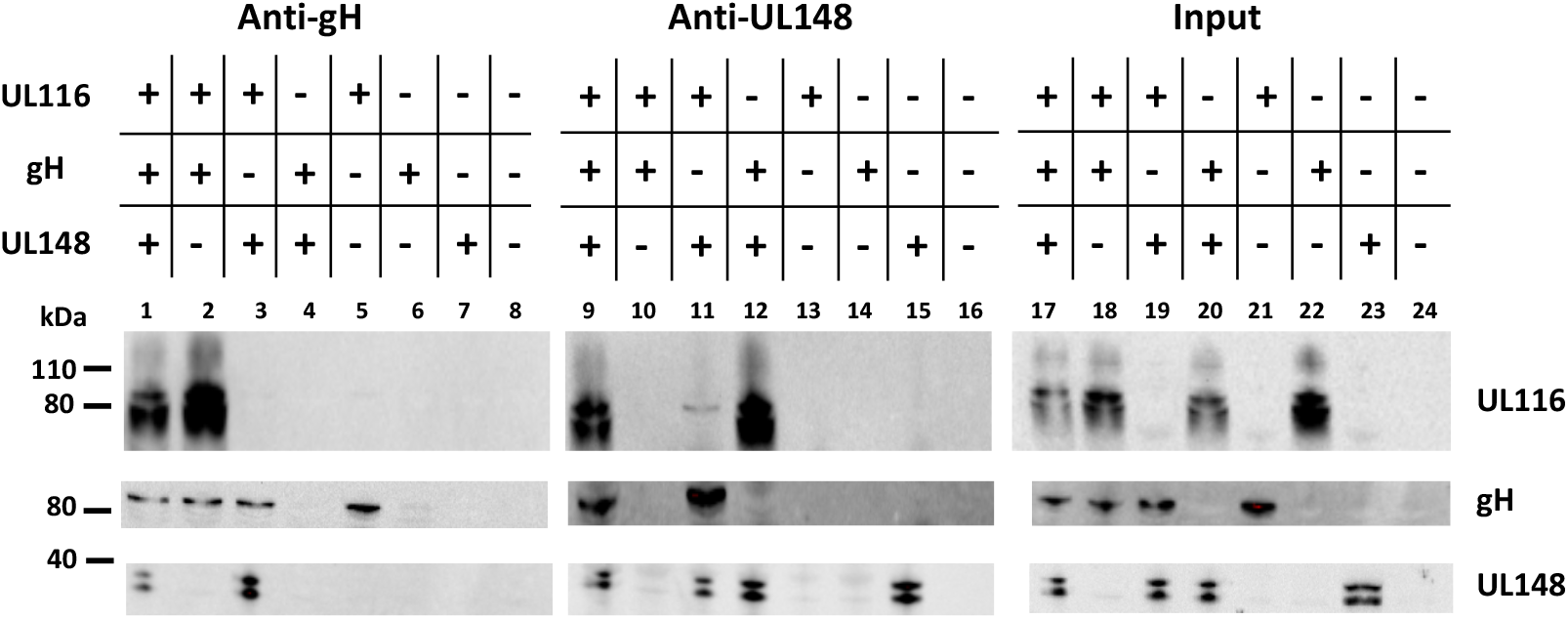
Co-immunoprecipitaions from transfected cells. pCDNA3.1(-)-gH_myc, pCDNA3.1(-)-UL116_strep and pCDNA3.1(-)-UL148_mycHis were used to transiently transfect HEK-293T cells, either alone or in combination to each other as specified on the top of the figure. pCDNA3.1(−) was used as control (lanes 8, 16 and 24). 48 hrs after transfection, cells were collected and the cleared lysates split in two aliquots for immunoprecipitation with anti-gH (human monoclonal MSL-109 antibody) and anti-UL148 (rabbit anti-His). UL-116 was revealed by the mouse monoclonal F11 antibody while anti-myc (mouse monoclonal) was used as probe to reveal gH-myc and UL148_mycHis. The western blot is representative of three independent experiments.

All together, these results are consistent with a direct interaction between UL116 and UL148, indicating a possible coordination of these two proteins in the ER for the determination of gH-based complexes formation.

## DISCUSSION

In our previous study, we showed that the HCMV UL116 protein is a non-disulfide bound gH-associated factor alternative to gL and that the complex is inserted into the viral envelope of mature particles (21). We sought to further characterize the role of UL116 in the HCMV life cycle by generating UL116-null virus and checking the cell-free infectivity of the progeny. Consistent with the current literature (31, 32), we found that UL116 in a nonessential protein and the TRG-*UL116*-null mutant virus was able to infect both fibroblasts and epithelial cells although producing roughly 10- and 6-fold less virus respectively. The TRG used in this study showed a roughly identical cell-free replication in cultured fibroblasts and epithelial cells as well as the TRG-*UL148*-null mutant. This last results is in contrast with a previous report from the literature where the TB40-*UL148*-null virus increases replication in epithelial cells (19). As it was pointed out in a very recent and elegant report, we suppose that this difference may be due to the genetic background of different strains (33). As first, we speculated that the gH/UL116 dimer could recognize a receptor on target cells surface and we purified the complex and checked binding by FACS. The data we obtained indicate the absence of a high affinity binding and we did not further check other possibilities but proceeded focusing on the intracellular role of UL116.

The TRG wt virus used in this study showed predominant Trimer over Pentamer on virions, consistent with previous reports (12). The relative amount of both complexes was strongly reduced in purified viruses from the recombinant TRG-*UL116*-null mutant likely not as defect in synthesis but rather as impaired incorporation in infectious virions. We found that gH levels in infected cells are partially rescued following treatment with an ERAD inhibitor indicating that UL116 does act on gH turn over and likely on its correct folding. gH-based complexes in infected cells show a milder reduction, once more suggesting that the impaired step rely on the efficiency of the complexes’ assembly. Indeed, virions derived from the TRG-*UL148*-null mutant were defective in the incorporation of Trimer but showed higher levels of Pentamer according to what reported in literature (19). Differently from ULl48, known to favor Trimer formation versus Pentamer, lack of UL116 impair both gH/gL derived complexes thus its action must be rather on proper assistance to gH. These findings suggest that UL116 is part of the molecular machinery required for the correct maturation and assembly of the complexes.

The lower amount of Trimer and Pentamer on viral particles reduces but does not abolish cell-free infectivity of the virus, the reason why UL116 was not recognized as an essential protein (31, 32). The group led by Dr. Ryckman has performed a deep analysis on the relationship between the major HCMV envelope glycoproteins and viral infectivity. Among others, their reports showed that the cell-free infectivity is modulated by the relative ratio of Trimer and Pentamer incorporated into the virion and that, although Pentamer definitely control epithelial tropism, its abundance is not straightforward correlated with efficiency of infection in non-fibroblast cell types (9, 12, 18). Trimer alone is enough for entry into fibroblasts (13) whereas Pentamer, always required for infection in all other cell types (12, 13), extends viral tropism through recognition of specific receptors recently identified (16, 17). Although the cell type restricted receptors explain the tropism specificity, the molecular mechanism responsible for viral infectivity depends several factors including glycoprotein isoforms, relative ratio of the complexes, RL13 locus and still non identified loci (9, 12, 18, 34, 35). Our findings show that cell-free infectivity is slightly modified but still functional at reduced levels of gH-based complexes on viral particles and address the mechanism of the molecular machinery regulating HCMV glycoproteins assembly.

The choice of gH/gL complexes carried by the mature virions starts during the early phases of glycoproteins assembly in the ER. To date, two viral proteins favoring formation of either Trimer or Pentamer have been identified: UL148 and US16 respectively. The single transmembrane (TM) spanning ER resident UL148 protein promotes gO incorporation without interacting with this glycoprotein but rather subtracting it to degradation by specifically targeting the ERAD receptor Sel1L (19, 36). This interaction activates unfolding protein response (UPR) leading to an ER expansion whose benefit for viral replication remains unclear (37). US16 is a 7TM HCMV protein identified as tropism factor whose absence impairs viral replication in epithelial/endothelial cells at the level of entry or post-entry (38). Remarkably, US16 is required for incorporation of UL128-131 showing a direct interaction only with UL130 (20). UL148 and US16 favor the incorporation of either gO or ULs respectively, harmonizing the correct formation of envelope complexes and highlighting that the broad tropism is due to a fine regulation of complexes levels.

From these data, we propose the following model of viral proteins interactions during the early ER post-synthesis phase (figure 8). We hypothesize that UL116 is the first interactor of gH, stabilizing the protein and protecting from degradation. To note, that gL (*UL115*) and *UL116* are adjacent genes on the same transcription unit and that the two proteins are bona fide synthesized simultaneously. As second step, UL148 interacts potentially with both gH and UL116 mediating binding to gL, a process that in absence of UL116 occurs at lower yield. It is possible that the formation of the gH/UL116 dimers progress while UL148 is involved in interaction with other factors such as the ERAD component Sel1L (36) while in its absence gH/UL116 dimer formation is favored. In absence of UL148, the heterodimer gH/UL116 is more stable and reduce the formation of gH/gL accessible for Trimer but especially for Pentamer assemble. Two disulfide bonds lock gL to gH and an additional cysteine on gL establish an alternative disulfide bridge to gO or UL128/UL130/UL131A resulting into the trimeric or pentameric complex respectively (22, 39). Thus, the noncovalent nature of gH/UL116 binding is ideal to chaperon gH toward a native or near-native conformation inducing stable conformer able to avoid host ERAD but also to be conformational competent to bind gL. Conformational instability of gH in absence of UL116 would explain why the levels of both the Trimer and Pentamer were lowered in *UL116*-null virions. UL148 would act downstream of UL116 as a regulatory factor acting on gH to favor gO incorporation on gH/gL. The role of US16 could be to stabilize the UL128/130/131A making this trimer available for incorporation. Although possible, any interaction between UL148 and US16 remains hypothetical. A further level of regulation can be hypothesized looking at the topology of US16 from Merlin (uniport Q6SVZ0). The cytoplasmatic C-terminal of US16 has 43 residues with 9 serines and 3 threonines that are putative sites of phosphorylation. Apart for the numerous kinases that could mediate ser/thr phosphorylation, the UL148 protein has been shown to activate the unfolding protein response (UPR) including the protein kinase R (PKR)-like ER kinase (PERK). PERK is a ser/thr kinase acting on restricted substrates (40) that is known to mediate efficient HCMV replication (41). Although this possibility is purely hypothetical, other 7TM protein of the early secretory pathway have been found to be regulated by cytoplasmic tail phosphorylation. For example, serine phosphorylation of the KDEL receptor by protein kinase A (PKA) is a crucial step in the regulation of the retrograde trafficking from Golgi to ER-Golgi (42). Intriguingly, the HCMV US17 gene product, known as interfering with the host innate immunoresponse, seems to play a role in controlling the viral level of gH. Guerzsinky et al. have shown that the reconstituted AD169 knocked out of the US17 gene show about 3 times reduction of viral gH without impairment of fibroblasts infectivity (43). In common with UL148, US17 interferes with the ER stress response inducing aberrant expression of several genes of this pathway. However, no data are available on a direct binding of this protein to gH and the observed reduction of gH levels in the US17 knock-out mutant could be an indirect effect such as an altered trafficking to the assembly complex (43).

**Figure 8.**
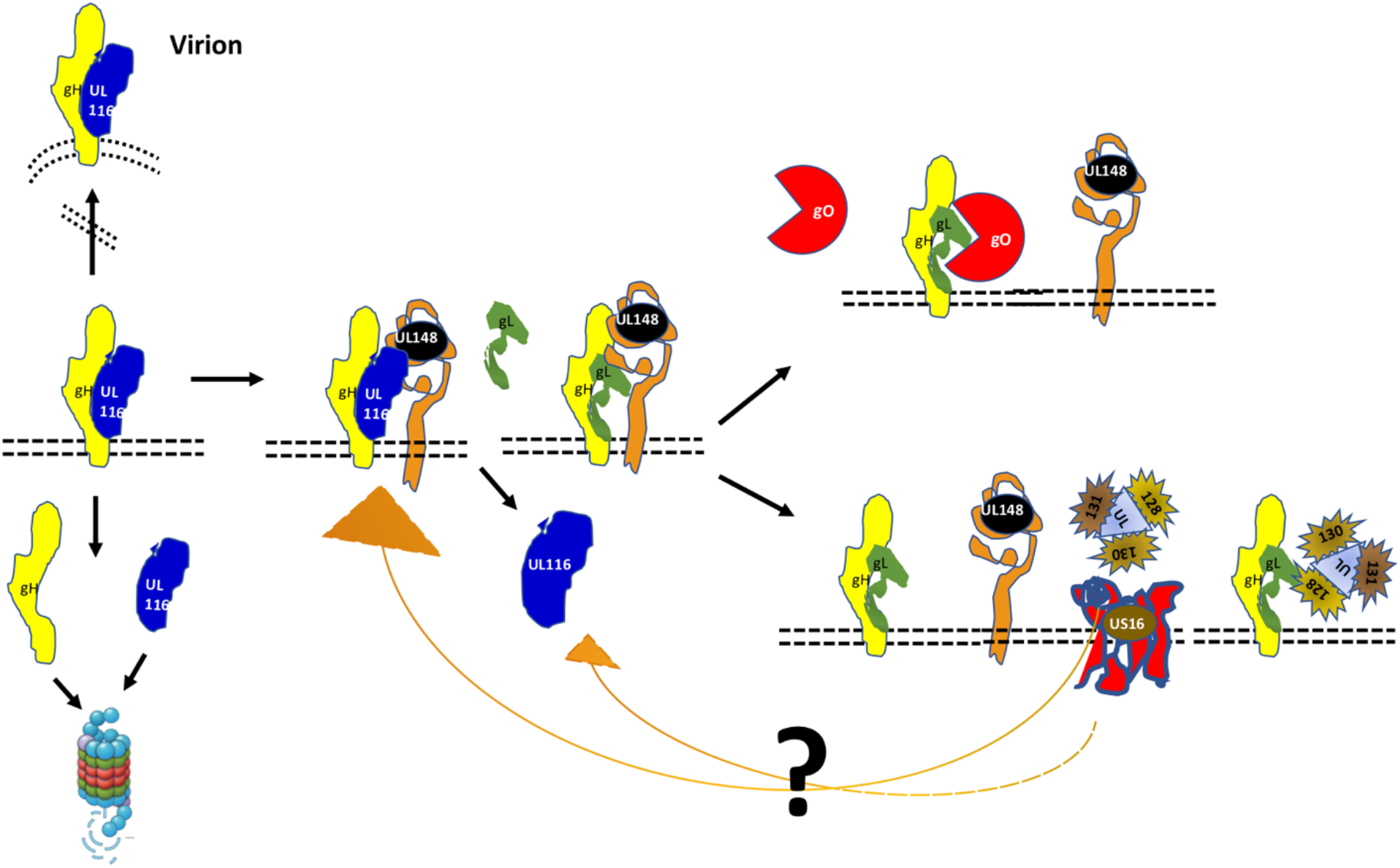
Model of HCMV interaction in the early phases of gH complexes assembly. UL116 is the first interactor of gH and chaperones the early folding steps. UL148 is recruited through either gH or UL116 and favors the binding of gL and successive association of gO. At limiting availability of UL148, for example engaged by Sel1A, UL116 remains bound to gH and traffic through the secretory pathway reaching the assembly complex and then the mature virion. UL116 can also be released from gH in absence of UL148, either at low efficiency or by the intervention of US16 or an unknown factor, allowing gL binding and favoring further incorporation of UL128-131 versus gO. Interaction of US16 with the other HCMV proteins is merely speculative.

In common with UL148 and US16, UL116 is also a nonessential viral protein (31, 32), highly conserved among strains that may suggest multiple interaction with other host and viral proteins (44). Our findings demonstrate that UL116 is required for reaching wt levels of both gH-based complexes but more generally of the viral particles’ levels of gH. Remarkably, in the *UL116*-null virus non disulfide bond gH in viral particles was completely missing and intracellular amount drastically reduced likely due to accelerate gH degradation. The presence of noncovalently linked gH was firstly revealed by Britt and collaborators while they identified a gp125 glycoprotein, then named gO, as part of gH/gL complex (45). Additionally, early characterization of the Pentamer by Wang and Shenk, preliminary described as gH/gL/UL128/UL130 complex, revealed a huge amount of noncovalently linked gH in infected cells (46). In this work we show that this fraction corresponds to gH associated to UL116 that, in addition, roughly represent the major gH complex carried by virions at least in TR. Indeed, in absence of UL148, the amount of gH/UL116 dimer further increases on viral particles as well as intracellularly. The direct interaction between UL148 and UL116 shown here suggests that the two proteins compete for gH association and that the formation of the disulfide bonds with gL is induced by UL148. Altogether, UL116 is a newly identified player of the molecular machinery responsible for the efficient folding and incorporation of gH-based complexes into virions

As UL116 protects gH from degradation in the ER, its meets the definition of molecular chaperone/escort as “any protein that interacts with and aids in the folding or assembly of another protein” and increase the yield of its client(s) (47). It would be also interesting to define the association of HCMV glycoproteins and folding assisting factors with ER cellular chaperones and/or other host proteins. A difference in the cell type host factors interacting with this machinery could explain why the levels of cellular pools of complexes are different between HFF and ARPE-19 cells. The best documented example of host contribution to the switch of envelope glycoproteins composition comes from the gammaherpesivirus EBV the infection of which is mainly restricted to B and epithelial cells. The gH/gL accessory protein gp42 is retained into the ER by the class II HLA, its receptor on B-cells, generating a progeny of gH/gL carrying virions with epithelial cells tropism (reviewed in (48)). In this case, the same host protein acts as receptor on B-cells for viral entry and as ER retaining factor for the viral ligand in the same cell type. For HCMV structural proteins such kind of analysis has not been performed yet. However, a molecular chaperone is not part of the final complex (47) while UL116 heterodimerizes with gH and is found on the viral envelope (21). This virion complex is massively represented on virions and could have irrelevant functions for HCMV pathogenesis as well as perform a still unnoticed role. The reduction of about 10-times cell-free viral infection in fibroblasts was reminiscent of adhesion factors from other *herpesviridae*. For instance, HSV gC protein acts as a “tethering” factor targeting glycosaminoglycans (GAGs) on cells (49) or the very abundant EBV gp350/220 glycoprotein that binds CD35 on host cells (50). These viral proteins are nonessential for entry, but they increase about 10-times viral cell-free infectivity. The experiments we have performed was not suitable to reveal eventual low affinity binding and we cannot exclude that a similar function belongs to gH/UL116. Further studies are required to dissect the function role of envelope UL116.

## Acknowledgments

We are in debt with Dr. Jeremy Kamil (LSU) for the fruitful discussion on UL148 and UL116 and Dr. Adam Feire (NIBR) for the generous gift of anti-gH MSL-109 antibody. We thank Simona Tavarini and Chiara Sammicheli (GSK) for FACS assistance.

## Contributorship

DY, SC, DM, and MM were involved in the conception and design of the study. GV, DA acquired the data. GV, DA, EF, DM, and MM analyzed and interpreted the results. All authors were involved in drafting the manuscript or revising it critically for important intellectual content. All authors had full access to the data and approved the manuscript before it was submitted by the corresponding author.

## REFERENCES

1. Mocarski ES, Shenk T, Pass RF. 2013. Cytomegaloviruses. In: Knipe DM, Howley PM, Fields Virology. Wolters Kluwer, Lippincott Williams & Wilkins.

2. Collins-McMillen D, Buehler J, Peppenelli M, Goodrum F. 2018. Molecular Determinants and the Regulation of Human Cytomegalovirus Latency and Reactivation. Viruses 10.

3. Zhuravskaya T, Maciejewski JP, Netski DM, Bruening E, Mackintosh FR, St Jeor S. 1997. Spread of human cytomegalovirus (HCMV) after infection of human hematopoietic progenitor cells: model of HCMV latency. Blood 90:2482–91.

4. Styczynski J. 2018. Who Is the Patient at Risk of CMV Recurrence: A Review of the Current Scientific Evidence with a Focus on Hematopoietic Cell Transplantation. Infect Dis Ther 7:1–16.

5. Streblow DN, Dumortier J, Moses AV, Orloff SL, Nelson JA. 2008. Mechanisms of cytomegalovirus-accelerated vascular disease: induction of paracrine factors that promote angiogenesis and wound healing. Curr Top Microbiol Immunol 325:397–415.

6. Boppana SB, Ross SA, Fowler KB. 2013. Congenital cytomegalovirus infection: clinical outcome. Clin Infect Dis 57 Suppl 4:S178–81.

7. Sinzger C, Digel M, Jahn G. 2008. Cytomegalovirus cell tropism. Curr Top Microbiol Immunol 325:63–83.

8. Heldwein EE, Krummenacher C. 2008. Entry of herpesviruses into mammalian cells. Cell Mol Life Sci 65:1653–68.

9. Zhou M, Yu Q, Wechsler A, Ryckman BJ. 2013. Comparative analysis of gO isoforms reveals that strains of human cytomegalovirus differ in the ratio of gH/gL/gO and gH/gL/UL128-131 in the virion envelope. J Virol 87:9680–90.

10. Ryckman BJ, Rainish BL, Chase MC, Borton JA, Nelson JA, Jarvis MA, Johnson DC. 2008. Characterization of the human cytomegalovirus gH/gL/UL128-131 complex that mediates entry into epithelial and endothelial cells. J Virol 82:60–70.

11. Huber MT, Compton T. 1997. Characterization of a novel third member of the human cytomegalovirus glycoprotein H-glycoprotein L complex. J Virol 71:5391–8.

12. Zhou M, Lanchy JM, Ryckman BJ. 2015. Human Cytomegalovirus gH/gL/gO Promotes the Fusion Step of Entry into All Cell Types, whereas gH/gL/UL128-131 Broadens Virus Tropism through a Distinct Mechanism. J Virol 89:8999–9009.

13. Kabanova A, Marcandalli J, Zhou T, Bianchi S, Baxa U, Tsybovsky Y, Lilleri D, Silacci-Fregni C, Foglierini M, Fernandez-Rodriguez BM, Druz A, Zhang B, Geiger R, Pagani M, Sallusto F, Kwong PD, Corti D, Lanzavecchia A, Perez L. 2016. Platelet-derived growth factor-alpha receptor is the cellular receptor for human cytomegalovirus gHgLgO trimer. Nat Microbiol 1:16082.

14. Wu Y, Prager A, Boos S, Resch M, Brizic I, Mach M, Wildner S, Scrivano L, Adler B. 2017. Human cytomegalovirus glycoprotein complex gH/gL/gO uses PDGFR-alpha as a key for entry. PLoS Pathog 13:e1006281.

15. Wu K, Oberstein A, Wang W, Shenk T. 2018. Role of PDGF receptor-alpha during human cytomegalovirus entry into fibroblasts. Proc Natl Acad Sci U S A 115:E9889–E9898.

16. Martinez-Martin N, Marcandalli J, Huang CS, Arthur CP, Perotti M, Foglierini M, Ho H, Dosey AM, Shriver S, Payandeh J, Leitner A, Lanzavecchia A, Perez L, Ciferri C. 2018. An Unbiased Screen for Human Cytomegalovirus Identifies Neuropilin-2 as a Central Viral Receptor. Cell 174:1158–1171 e19.

17. E X, Meraner P, Lu P, Perreira JM, Aker AM, McDougall WM, Zhuge R, Chan GC, Gerstein RM, Caposio P, Yurochko AD, Brass AL, Kowalik TF. 2019. OR14I1 is a receptor for the human cytomegalovirus pentameric complex and defines viral epithelial cell tropism. Proc Natl Acad Sci U S A 116:7043–7052.

18. Zhang L, Zhou M, Stanton R, Kamil J, Ryckman BJ. 2018. Expression Levels of Glycoprotein O (gO) Vary between Strains of Human Cytomegalovirus, Influencing the Assembly of gH/gL Complexes and Virion Infectivity. J Virol 92.

19. Li G, Nguyen CC, Ryckman BJ, Britt WJ, Kamil JP. 2015. A viral regulator of glycoprotein complexes contributes to human cytomegalovirus cell tropism. Proc Natl Acad Sci U S A 112:4471–6.

20. Luganini A, Cavaletto N, Raimondo S, Geuna S, Gribaudo G. 2017. Loss of the Human Cytomegalovirus US16 Protein Abrogates Virus Entry into Endothelial and Epithelial Cells by Reducing the Virion Content of the Pentamer. J Virol 91.

21. Calo S, Cortese M, Ciferri C, Bruno L, Gerrein R, Benucci B, Monda G, Gentile M, Kessler T, Uematsu Y, Maione D, Lilja AE, Carfi A, Merola M. 2016. The Human Cytomegalovirus UL116 Gene Encodes an Envelope Glycoprotein Forming a Complex with gH Independently from gL. J Virol 90:4926–38.

22. Ciferri C, Chandramouli S, Donnarumma D, Nikitin PA, Cianfrocco MA, Gerrein R, Feire AL, Barnett SW, Lilja AE, Rappuoli R, Norais N, Settembre EC, Carfi A. 2015. Structural and biochemical studies of HCMV gH/gL/gO and Pentamer reveal mutually exclusive cell entry complexes. Proc Natl Acad Sci U S A 112:1767–72.

23. Murphy E, Yu D, Grimwood J, Schmutz J, Dickson M, Jarvis MA, Hahn G, Nelson JA, Myers RM, Shenk TE. 2003. Coding potential of laboratory and clinical strains of human cytomegalovirus. Proc Natl Acad Sci U S A 100:14976–81.

24. Smith IL, Taskintuna I, Rahhal FM, Powell HC, Ai E, Mueller AJ, Spector SA, Freeman WR. 1998. Clinical failure of CMV retinitis with intravitreal cidofovir is associated with antiviral resistance. Arch Ophthalmol 116:178–85.

25. Britt W. 2010. Human Cytomegalovirus: Propagation, quantification and storage. Current Protocols in Microbiology 14E.3.

26. Tischer BK, von Einem J, Kaufer B, Osterrieder N. 2006. Two-step red-mediated recombination for versatile high-efficiency markerless DNA manipulation in Escherichia coli. Biotechniques 40:191–7.

27. Tischer BK, Smith GA, Osterrieder N. 2010. En passant mutagenesis: a two step markerless red recombination system. Methods Mol Biol 634:421–30.

28. Baldick CJ, Jr., Marchini A, Patterson CE, Shenk T. 1997. Human cytomegalovirus tegument protein pp71 (ppUL82) enhances the infectivity of viral DNA and accelerates the infectious cycle. J Virol 71:4400–8.

29. Tischer BK, Von Einem J, Kaufer B, Osterrieder N. 2006. Two-step Red-mediated recombination for versatile high-efficiency markerless DNA manipulation in Escherichia coli. Biotechniques 40:191–197.

30. Wang F, Song W, Brancati G, Segatori L. 2011. Inhibition of endoplasmic reticulum-associated degradation rescues native folding in loss of function protein misfolding diseases. J Biol Chem 286:43454–64.

31. Yu D, Silva MC, Shenk T. 2003. Functional map of human cytomegalovirus AD169 defined by global mutational analysis. Proc Natl Acad Sci U S A 100:12396–401.

32. Dunn W, Chou C, Li H, Hai R, Patterson D, Stolc V, Zhu H, Liu F. 2003. Functional profiling of a human cytomegalovirus genome. Proc Natl Acad Sci U S A 100:14223–8.

33. Day LZ, Stegmann C, Schultz EP, Lanchy JM, Yu Q, Ryckman BJ. 2020. Polymorphisms in Human Cytomegalovirus gO Exert Epistatic Influences on Cell-Free and Cell-To-Cell Spread, and Antibody Neutralization on gH Epitopes. J Virol doi:10.1128/JVI.02051-19.

34. Stanton RJ, Baluchova K, Dargan DJ, Cunningham C, Sheehy O, Seirafian S, McSharry BP, Neale ML, Davies JA, Tomasec P, Davison AJ, Wilkinson GWG. 2010. Reconstruction of the complete human cytomegalovirus genome in a BAC reveals RL13 to be a potent inhibitor of replication. The Journal of Clinical Investigation 120:3191–3208.

35. Schultz EP, Lanchy JM, Day LZ, Yu Q, Peterson C, Preece J, Ryckman BJ. 2020. Specialization for Cell-Free or Cell-to-Cell Spread of BAC-Cloned Human Cytomegalovirus Strains Is Determined by Factors beyond the UL128-131 and RL13 Loci. J Virol 94.

36. Nguyen CC, Siddiquey MNA, Zhang H, Li G, Kamil JP. 2018. Human Cytomegalovirus Tropism Modulator UL148 Interacts with SEL1L, a Cellular Factor That Governs Endoplasmic Reticulum-Associated Degradation of the Viral Envelope Glycoprotein gO. J Virol 92.

37. Siddiquey MNA, Zhang H, Nguyen CC, Domma AJ, Kamil JP. 2018. The Human Cytomegalovirus Endoplasmic Reticulum-Resident Glycoprotein UL148 Activates the Unfolded Protein Response. J Virol 92.

38. Bronzini M, Luganini A, Dell’Oste V, De Andrea M, Landolfo S, Gribaudo G. 2012. The US16 gene of human cytomegalovirus is required for efficient viral infection of endothelial and epithelial cells. J Virol 86:6875–88.

39. Chandramouli S, Malito E, Nguyen T, Luisi K, Donnarumma D, Xing Y, Norais N, Yu D, Carfi A. 2017. Structural basis for potent antibody-mediated neutralization of human cytomegalovirus. Sci Immunol 2.

40. Pytel D, Majsterek I, Diehl JA. 2016. Tumor progression and the different faces of the PERK kinase. Oncogene 35:1207–15.

41. Yu Y, Pierciey FJ, Jr., Maguire TG, Alwine JC. 2013. PKR-like endoplasmic reticulum kinase is necessary for lipogenic activation during HCMV infection. PLoS Pathog 9:e1003266.

42. Cabrera M, Muniz M, Hidalgo J, Vega L, Martin ME, Velasco A. 2003. The retrieval function of the KDEL receptor requires PKA phosphorylation of its C-terminus. Mol Biol Cell 14:4114–25.

43. Gurczynski SJ, Das S, Pellett PE. 2014. Deletion of the human cytomegalovirus US17 gene increases the ratio of genomes per infectious unit and alters regulation of immune and endoplasmic reticulum stress response genes at early and late times after infection. J Virol 88:2168–82.

44. Foglierini M, Marcandalli J, Perez L. 2019. HCMV Envelope Glycoprotein Diversity Demystified. Front Microbiol 10:1005.

45. Li L, Nelson JA, Britt WJ. 1997. Glycoprotein H-related complexes of human cytomegalovirus: identification of a third protein in the gCIII complex. J Virol 71:3090–7.

46. Wang D, Shenk T. 2005. Human cytomegalovirus virion protein complex required for epithelial and endothelial cell tropism. Proc Natl Acad Sci U S A 102:18153–8.

47. Kim YE, Hipp MS, Bracher A, Hayer-Hartl M, Hartl FU. 2013. Molecular chaperone functions in protein folding and proteostasis. Annu Rev Biochem 82:323–55.

48. Mohl BS, Chen J, Sathiyamoorthy K, Jardetzky TS, Longnecker R. 2016. Structural and Mechanistic Insights into the Tropism of Epstein-Barr Virus. Mol Cells 39:286–91.

49. Herold BC, WuDunn D, Soltys N, Spear PG. 1991. Glycoprotein C of herpes simplex virus type 1 plays a principal role in the adsorption of virus to cells and in infectivity. J Virol 65:1090–8.

50. Ogembo JG, Kannan L, Ghiran I, Nicholson-Weller A, Finberg RW, Tsokos GC, Fingeroth JD. 2013. Human complement receptor type 1/CD35 is an Epstein-Barr Virus receptor. Cell Rep 3:371–85.

